# A new machine learning method for cancer mutation analysis

**DOI:** 10.1101/2022.06.29.498062

**Authors:** Mahnaz Habibi, Golnaz Taheri

## Abstract

It is complicated to identify cancer-causing mutations. The recurrence of a mutation in patients remains one of the most reliable features of mutation driver status. However, some mutations are more likely to happen than others for various reasons. Different sequencing analysis has revealed that cancer driver genes operate across complex pathways and networks, with mutations often arising in a mutually exclusive pattern. Genes with low-frequency mutations are understudied as cancer-related genes, especially in the context of networks. Here we propose a machine learning method to study the functionality of mutually exclusive genes in the networks derived from mutation associations, gene-gene interactions, and graph clustering. These networks have indicated critical biological components in the essential pathways, especially those mutated at low frequency. Studying the network and not just the impact of a single gene significantly increases the statistical power of clinical analysis. The proposed method identified important driver genes with different frequencies. We studied the function and the associated pathways in which the candidate driver genes participate. By introducing lower-frequency genes, we recognized less studied cancer-related pathways. We also proposed a novel clustering method to specify driver modules in each type of cancer. We evaluated each cluster with different criteria, including the terms of biological processes and the number of simultaneous mutations in each cancer. Materials and implementations are available at: https://github.com/MahnazHabibi/Mutation_Analysis

## 1 Introduction

The driving forces behind cancer are gene, nucleotide, and cellular structure changes. Somatic cells can acquire mutations one or two orders of magnitude more quickly than germline cells, making them more susceptible to different types of cancer [1]. The vast majority of these mutations, called passenger, have little effect on cell proliferation compared to a few driver mutations that give cells a selective advantage [2]. Mutations can activate or deactivate proteins, and they can change a wide range of cellular processes for different patients and types of cancer. This results in high intra-and inter-tumor heterogeneity in biochemistry and histology, which may explain why some cancers are resistant to treatment and make it more challenging to identify the events that cause cancer [3–5].

Next-generation sequencing technology has transformed the cancer genome study, facilitating us to sequence whole-genome and find somatic mutations in millions of cancer genomes. The Cancer Genome Atlas (TCGA), a publicly funded genomics project, contains a collection of mutation profiles from thousands of patients for more than 30 different types of cancer [6]. The recent mutation perspective demonstrates the importance of specifying genes and their associated networks to detect the cancer driver genes. Finding significantly mutated genes with high recurrent mutations can help us better predict the course of cancer development and progression. These cancer-causing driver genes are difficult to track down, and many of the mutations have not been detected using existing methods datasets. Methodical studies have shown multiple new genes and classes of cancer genes, respectively. They have also demonstrated that despite some cancer genes being mutated with high frequencies, most cancer genes in most patients arise with intermediate or low frequencies (2–20%). Therefore, a complete record of mutations in this frequency class will be essential for identifying dysregulated pathways and effective targets for therapeutic interference. Nevertheless, current studies present significant gaps in our understanding of cancer genes with intermediate frequency. For example, a study of 183 lung adenocarcinomas discovered that 15% of patients missed even a single mutation influencing any of the 10 known hallmarks of cancer, and 38% of patients had 3 or even fewer such mutations [7]. As a result, we cannot capture a complete expression profile of all genes and subsets of genes that drive the evolution and progression of cancer. Cancer genes tend to alter considerably in a limited number of pathways, especially in pathways related to survival, cell division, differentiation, and genomic preservation. Therefore, it is necessary to determine the pathway-level importance of genes, even those genes mutated at low frequencies [8].

Since mutually exclusive couples of genes usually share similar pathways, one strategy for detecting these drivers is to explore for mutual exclusivity of changed genes. However, we know that mutated genes seldom coexist in the same tumor, while only one gene in a pathway is typically found to have a driver mutation in each patient [9]. This situation may occur due to cancer pathways’ functional redundancy or synthetic lethality. Typical examples of mutually exclusive driver mutations contain EGFR and KRAS mutations in lung cancer [10] and TP53 and MDM2 mutations in glioblastoma [6]. Based on this explanation, finding mutual exclusivity modules in cancer needs to find important and more relevant genes, find the correlation between them, and analyze them. Then this analysis needs statistical tests to identify network modules demonstrating patterns of mutually exclusive genetic changes across multiple patients [11]. As a new method, Mutex uses a large pathway model of human signaling processes to explore groups of mutually exclusively changed genes that share a joint downstream event [12].

The main disadvantage of the current methods is that they need comprehensive filtering of mutation data, which are restricted to the most significantly mutated genes and concentrate on predefined network modules [13]. The mutual exclusivity signal may be biased towards recognizing gene sets where most of the coverage comes just from highly mutated genes [14, 15]. Although cancer-related genes have been shown to be involved in numerous pathways, few methods determine the important gene sets where a gene has various mutually exclusive correlations with other genes in diverse pathways at different mutation frequencies. We proposed a novel three-step method to identify candidate driver gene sets with mutually exclusive mutations to find the mutually exclusive mutation pattern more comprehensively. In the first step, the proposed unsupervised machine learning method detects candidate driver genes. For this purpose, we constructed a biological network corresponding to important cancer-related genes. Then, we defined six informative topological and biological features for each gene as a node in the network. We calculated the score for our predefined features for each gene. Afterward, we introduced the high-score genes with meaningful relationships to cancer as candidates for more investigation. In the second step, we presented a network based on the relationship between genes to identify the cancer-related clusters. We used the information on physical, biological, and functional interaction between the high-score candidate driver genes obtained in the first step to construct this network. In the last step, we specify driver modules of different sizes with various cutoffs and importance and the number of simultaneous mutations in each cancer, from the previous step clusters.

## 2 Materials and methods

In this section, we present a new three-step method for identifying driver genes and modules in different types of cancer. In the first step, we proposed an unsupervised machine learning method to recognize a set of candidate driver mutated genes associated with different types of cancer. In this step, we used the information of different patients (cases) with various types of cancer and their associated mutated genes to create a weighted network of mutated genes. Then six informative topological features are calculated for each gene as a node of the constructed network. We generated a feature matrix for the set of candidate mutated genes *X* = [*x*_*ij*_]_*m*×*n*_ that each *x*_*ij*_ component represents the *j*-th feature for the *i*-th gene. Then we employed an unsupervised learning method to calculate the appropriate scores for each of the predefined features. Finally, our proposed method selects a set of genes with higher scores as a set of mutated genes that contain valuable information. In the second step, we constructed a network based on the relationship between genes to identify the cancer-related clusters. We used the information on biological and functional interactions between the high-score genes obtained in the first step to build this network. Then, we used a heuristic method to cluster the constructed weighted network. The weight of each cluster is calculated based on the average weight of the nodes of that cluster. The set of clusters with higher weights is identified as the cancer-related clusters containing important information. In the last step, in each cluster, we identified driver modules for each cancer based on the number of cases in which the specific gene was mutated in different types of cancer. The general workflow of the proposed method is illustrated in Figure 1.

**Fig 1.**
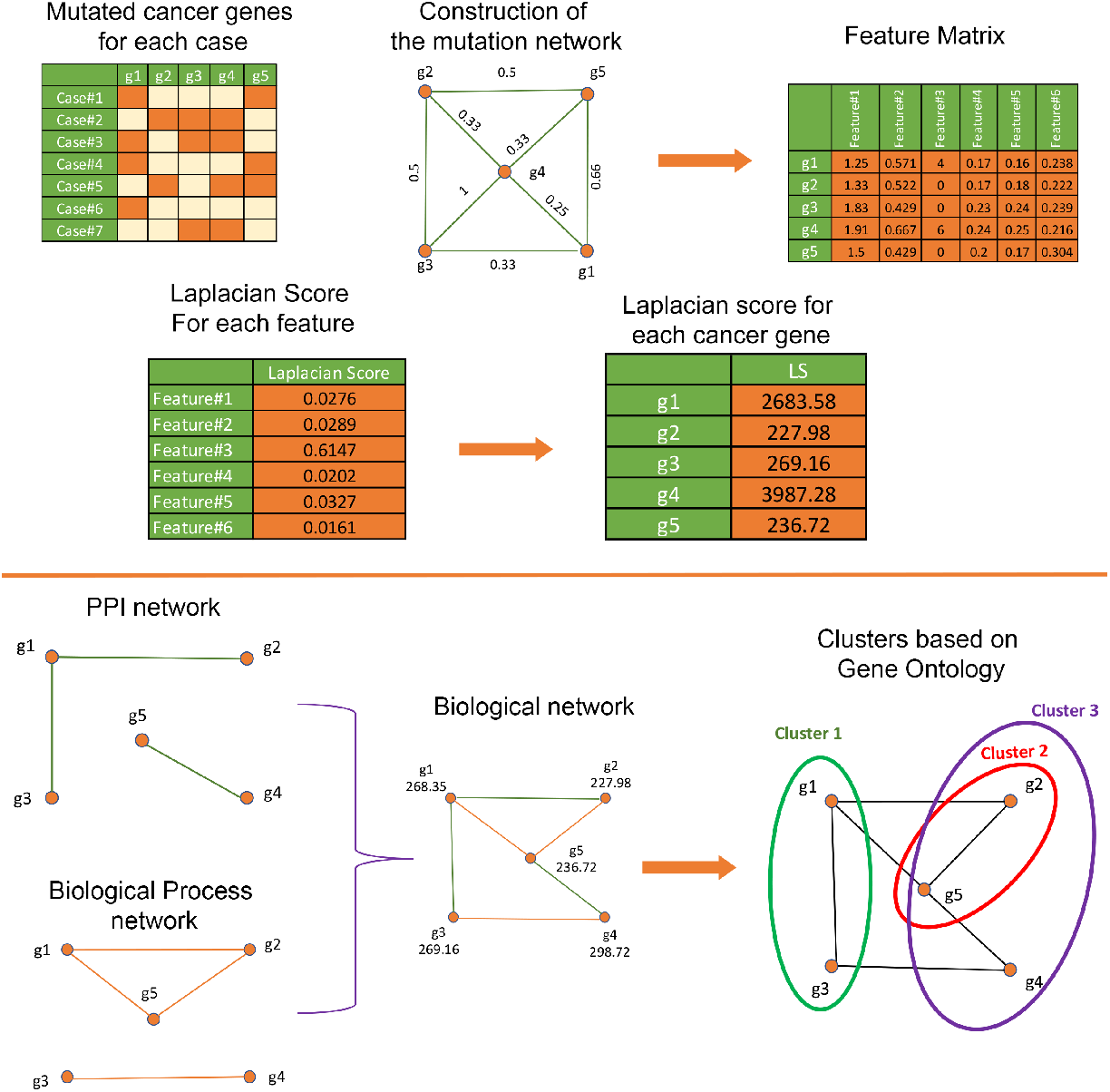
The workflow of the proposed method.

### 2.1 Datasets

Identifying associated driver genes with different cancer types plays a significant role in determining mutated driver modules. Therefore, the starting point is identifying appropriate datasets to extract complete information about the somatic mutation, corresponding protein-protein interaction (PPI), and biological process information. A representative set of tumors and mutations were gathered from TCGA, on March 2022 [6]. We downloaded the information on the primary site and mutations for 12,792 cases. This dataset contains 576 mutated cancer genes and 15 major primary sites. We used the PPI network from Habibi et al. (2021) [16]. This dataset contains the physical interactions between proteins that are collected from the Biological General Repository for Interaction Datasets (BioGRID) [17], Agile Protein Interactomes Data analyzer (APID) [18], Homologous interactions (Hint) [19], Human Integrated Protein-Protein Interaction reference (HIPPIE) [20] and Huri [21]. All of the proteins in this dataset are mapped to a universal protein resource (UniProt) ID [22]. This interactome contains 20,040 proteins and 304,730 interactions. We also used the informative biological processes related to each mutated gene that is gathered from the Gene Ontology (GO) [23], to identify functional interactions between mutated cancer genes.

### 2.2 Construction of the mutation network

We introduced a mutation network based on 576 mutated cancer genes in this work. Suppose that *V* = {*g*_1_, …, *g*_*n*_} indicates the set of mutated cancer genes. Also, suppose that 𝒸(*g*_*i*_) is the set of cases that contain a given mutation gene (*g*_*i*_). A weighted mutation network *G* =*< V, E, ω >* was constructed by connecting two genes *g*_*i*_ and *g*_*j*_ if and only if 𝒸(*g*_*i*_) ∩ 𝒸(*g*_*j*_) ≠ ∅. The weight of edge *g*_*i*_*g*_*j*_ ∈ *E* which is denoted by *ω*(*g*_*i*_*g*_*j*_), is defined as follows:

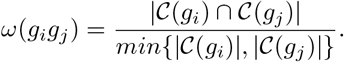

A path between *g*_*i*_ and *g*_*j*_ is determined as a sequence of distinct nodes such that an edge of *G* connects two consequent nodes. The weight of a path equals the sum of the weights of edges in this path. The shortest path from node *g*_*i*_ to node *g*_*j*_ is a path between two nodes with minimum weight. The weight of the shortest path between two nodes *g*_*i*_ and *g*_*j*_ is denoted by *d*_*w*_(*g*_*i*_, *g*_*j*_).

#### 2.2.1 Informative topological features for mutation network

We defined the following informative topological features for each node of the weighed mutated network.

- **Weight**: The Weight of node *g*_*i*_ on weighted graph *G* =*< V, E, ω >* as follows:

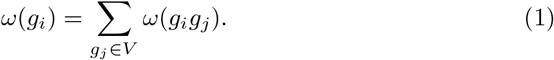
- **Closeness**: The Closeness centrality measure is defined for each node, *g*_*i*_, as follows:

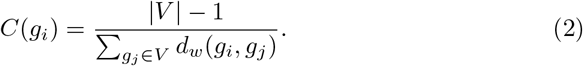
- **Betweenness**: The Betweenness centrality measure is defined of each node *g*_*i*_ on network *G* as follows:

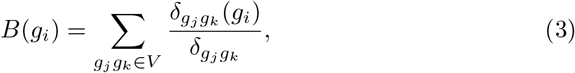

where*δ*_*gj gk*_ denoted the weights of shortest paths between two nodes *g*_*i*_ and *g*_*k*_ and *δ*_*gjgk*_ (*g*_*i*_) is indicated the weighs of shortest paths between two nodes *g*_*i*_ and *g*_*k*_ pass through node *g*_*i*_.
- **PageRank**: The score for each node *g*_*i*_ in the network is calculated based on all the scores assigned to all nodes *g*_*j*_, which are connected iteratively as follows:

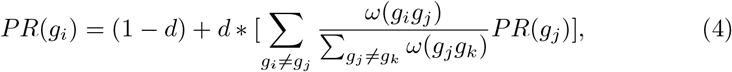

where *d* is a parameter between 0 and 1.
- **Eigenvector**: The Eigenvector centrality measure is defined as the amount of influence for a node *g*_*i*_ in the network as follows:

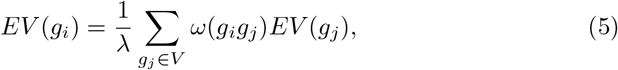

where *λ* is a constant.
- **Entropy**: Suppose that *ω*(*g*_*i*_) is the weight of node *g*_*i*_ on weighted network *G* =*< V, E, ω >*. The probability distribution vector ? =*< π*_1_, …, *π* _(|*V* |)_ *>* is defined on set of all nodes of the network as follows:

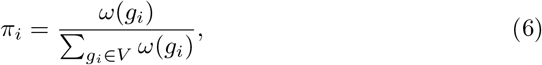

Then, the entropy of weighted graph *G* is calculated as follows:

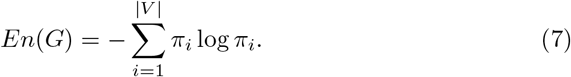

We calculated the effect of each node *g*_*i*_ on network entropy as follows:

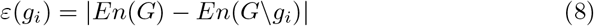

where *G*\*g*_*i*_ is the weighted network that is constructed with respect to the removal of node *g*_*i*_ and its connected edges from the network.

### 2.3 Machine learning method to select top mutated cancer genes

Since the problem of selecting the set of mutated candidate driver cancer genes is still an open question, it can be studied as a problem without an exact answer. Therefore, we utilized an effective unsupervised feature selection method to determine an efficient set of mutated cancer genes. Suppose that *X* = [*x*_*ij*_]_*m*×*n*_ represents the feature matrix and *x*_*ij*_ represents the *j*-th feature of the *i*-th sample (genes). We assigned a feature vector 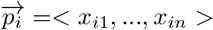 to each sample and defined the column matrix *F*_*j*_ = [*x*_1*j*_, …, *x*_*mj*_]^*T*^ for the *j*-th feature. To find an appreciated score for each feature, we used the Laplacian Score for Feature Selection (LSFS) as an unsupervised machine learning method as follows:

Suppose that *S* = [*s*_*ij*_]_*m*×*m*_ indicates the weighted matrix where 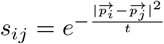 *t* if the euclidean distance between two feature vectors 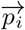and 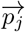 is less than *δ*. Also, *k*=1 suppose that *D* = [*d*_*i*_]_*m*×*m*_ is the diagonal matrix where 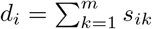 and *L* = *D* – *S* is the Laplacian matrix. The Laplacian Score for each feature, *j*, is calculated as follows:

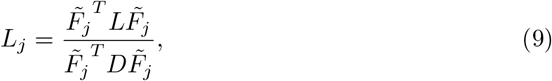

where *J* = [1, 1,.., 1]^*T*^ and 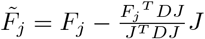.

Finally, we calculated the LS for each mutated cancer gene *g*_*i*_ as follows:

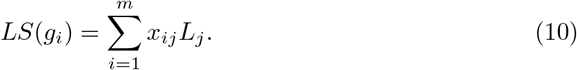

The algorithm to calculate Laplacian Score (LS) for each mutated cancer gene is described in Algorithm 1.

### 2.4 Heuristic algorithm to identify specific modules for each cancer type

A biological network is constructed as an undirect weighted graph 𝒢 =*<* 𝒱, ℰ, 𝒲; *>* where the set of nodes 𝒱= {*g*_1_, …, *g*_*N*_ } is the *N* top mutated cancer genes regarding maximum LS values. Two mutated cancer genes *g*_*i*_ and *g*_*j*_ are connected through an edge *e*_*ij*_ if they participate in the same biological process or if there is physical interaction between them. The 𝒲(*g*_*i*_) represents the weight of the mutated cancer gene with respect LS value. In the following, we present a heuristic algorithm *MG* to cluster the weighted network 𝒢. Suppose that 𝒮 ⊆ 𝒱 is the subset of nodes in the network. The neighborhood of 𝒮 is defined as follows:

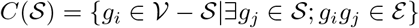

The *MG* algorithm first selects a node as a cluster. Then, the new cluster expands by adding a new node to the cluster regarding the average LS value of the nodes in this cluster and the LS values of adjacent nodes in the cluster. The *MG* algorithm adds a new adjacent node to a cluster such that the node’s weight is greater than the average weight of the cluster nodes. If the weight of all adjacent cluster nodes is less than the average cluster weight, the *MG* algorithm adds an adjacent node with the highest weight to the cluster with a small probability. The likelihood of reaching nodes with smaller weights decreases as the number of nodes in the cluster increases. The *MG* constructs a new cluster by selecting a new seed from the network nodes that have not been placed in a cluster and then expanding this node to get a new cluster with the maximum average weight of all nodes. The *MG* algorithm extends clusters to weighted graphs described in Algorithm 2.

#### Algorithm 1 The Laplacian Score (LS) algorithm

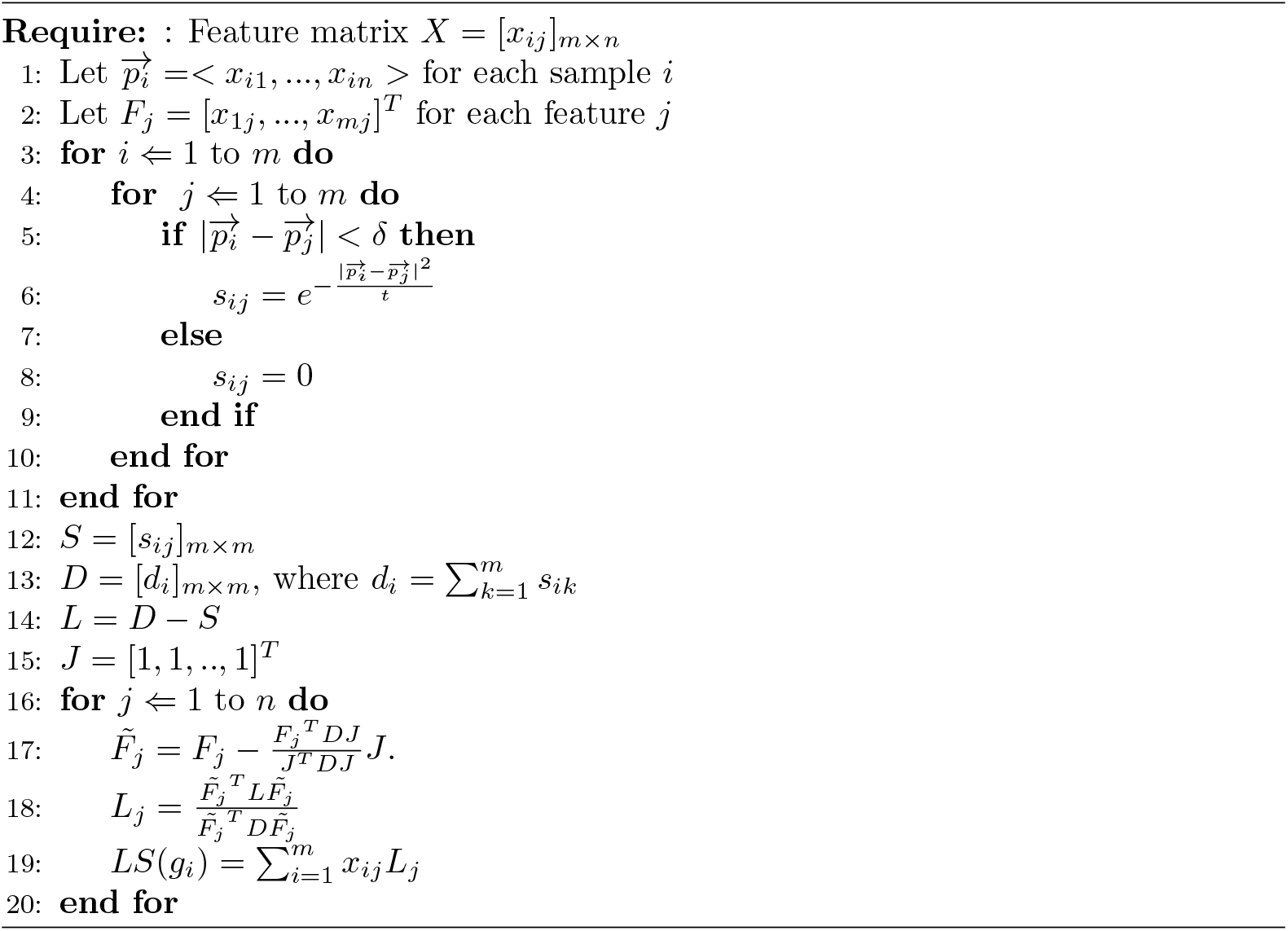

#### Algorithm 2 *MG* algorithm

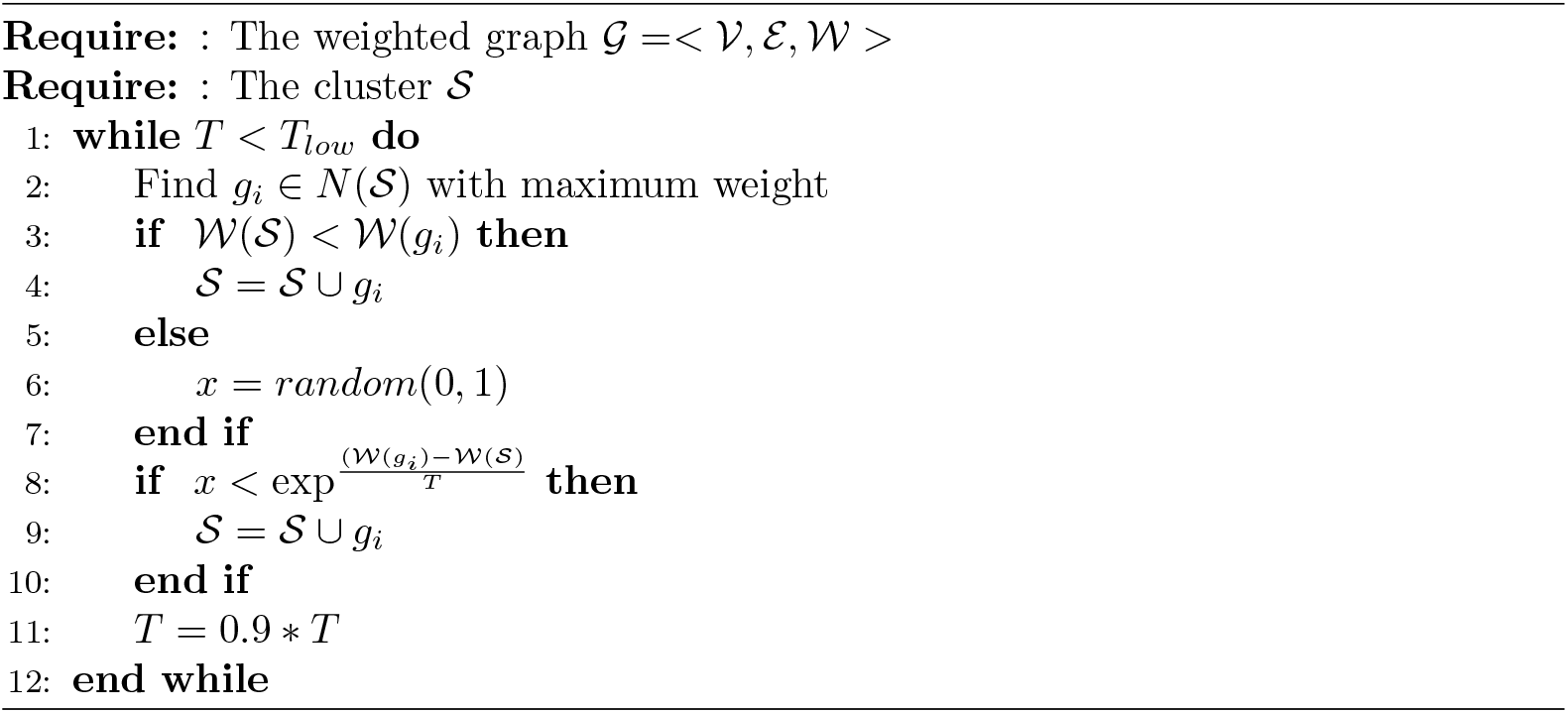

## 3 Results

### 3.1 Evaluation of high score selected genes based on Laplacian Score

One of the significant challenges for existing methods is that they need extensive filtering of mutation data, which is limited to the most significantly mutated genes and focuses on predefined modules. Therefore, the mutual exclusivity signal can be biased toward recognizing gene sets where most of the coverage comes from highly mutated genes. Finding a new set of genes with essential properties, even if they have moderately or infrequently mutated, leads us to some new informative modules. The proposed unsupervised machine learning method selects a list of 200 mutated genes with high scores through its predefined properties. Figure 2 shows the heat map of the number of mutations for each gene in each cancer and the value of the associated LS for these high-score genes. In Figure 2, we sorted 200 high-score genes based on the number of their mutations in 15 different types of cancer. Genes with high LS are highlighted in Figure 2. This figure contains some of the frequently mutated genes such as TP53, FAT4, and KMT2C, and some of the infrequently mutated genes such as FSTL3, SSX2, and MDS2. Since most recent studies have focused on frequently mutated genes, we also studied the number of infrequently mutated genes with high LS in addition to the frequently mutated genes. In the following, we present a list of these infrequently mutated genes with high LS.

**Fig 2.**
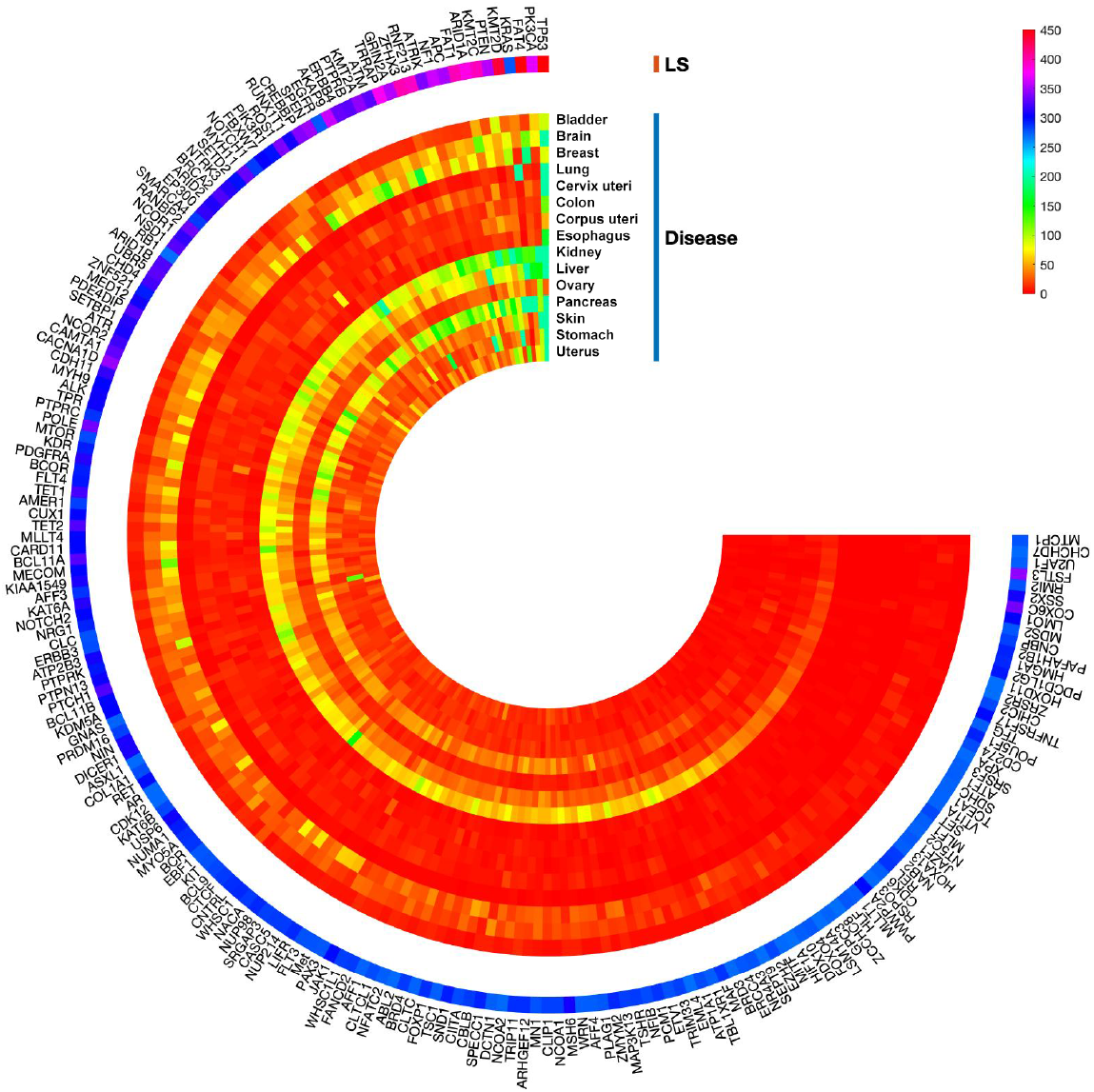
Cross-talk between mentioned signaling pathways in different types of cancer

- Follistatin-like 3 (FSTL3) is expressed in normal human tissues. Increasing evidence demonstrates that FSTL3 plays an essential role in regulating embryonic evolution, osteogenesis, glucose, and lipid metabolism. Furthermore, FSTL3 was found abundantly expressed in cell lung cancer and breast cancer and participates in tumor progression, containing invasion and metastasis. FSTL3 is an independent risk factor connected with the prognosis for different cancers [24].
- Recent studies demonstrate that Cytochrome c oxidase subunit 6c (COX6C) has a particular association with breast cancer, esophageal cancer, thyroid tumors, prostate cancer, uterine cancer, and melanoma. Several reports show that the differential expression of COX6C is associated with predicting some tumors and is expected to become one of the diagnostic markers of typical tumors [25].
- Recent studies demonstrated that SSX2 induces aging in different cells, as specified by classical aging features, including enlargement of the cytoplasm, cell growth arrest, and DNA double-strand breaks. SSX proteins are expressed in multiple types of tumors, such as 40% of melanomas and up to 65% of breast cancers. The SSX family comprises nine similar members, most likely redundant in their cellular functions [26].
- LMO1 belongs to the family of LIM-only domain genes (LMOs). Some studies have shown that LMO1 plays an essential role in the tumorigenesis of several types of cancer, including leukemia, breast cancer, and neuroblastoma. The author of [27] found that LMO1 was significantly over-expressed in non-small cell lung cancer (NSCLC) samples relative to normal adjacent tissue and that over-expression of LMO1 in NSCLC cells elevated cell proliferation, supporting an oncogenic function in NSCLC.
- The TNF receptor superfamily member 17 (TNFRSF17) is a gene that encodes a protein involved in B cell development and autoimmune response. This protein also plays a role in activating NF-*κ*B and MAPK8/JNK. Multiple types of mutations in TNFRSF17 have been shown in endometrial cancer, intestinal cancer, and skin cancer. On average, TNFRSF17 mutations are found in 0.50% of all cancers; the most common types are colorectal, colon cancer, glioblastoma, lung cancer, and malignant cancer melanomas [30].
- The Programmed cell death 1 ligand 2 (PD-L2) is a gene that encodes a protein that involves in the signal that is required for IFNG production and T-cell proliferation. Multiple types of mutations in PD-L2 have been observed in intestinal cancer, skin cancer, and stomach cancer. On average, PD-L2 mutations are found in 0.83% of all cancers; the most common types are lung cancer, breast invasive ductal carcinoma, colon cancer, urothelial bladder carcinoma, and high-grade ovarian cancer [29].
- POU5F1 is associated with the pluripotency and proliferative potential of ESCs and germ cells. Previous studies have shown that POU5F1 plays a critical role in maintaining the normal stem cell self-renewal process. Several studies have noted the expression of POUF1 in human cancer cells such as breast cancer, ovarian cancer, and melanoma. Moreover, recent studies revealed that POU5F1 expression was significantly elevated in tumor tissues compared to non-cancerous tissues [31].
- The high mobility group A1 (HMGA1) gene has an essential role in embryonic development. Multiple studies have shown elevated HMGA1 expression in malignant cancer such as breast cancer, lung cancer, colorectal cancer, and uterine cancer. Collectively, these studies reveal that HMGA1 has an essential role in tumorigenesis and tumor progression [32].
- The Programmed death-ligand 1 (PD-L1), also known as CD274 on cancer cells, contributes to cancer immune escape. The PD-1/PD-L1 axis is the major speed-limiting step of the anti-cancer immune response for multiple cancer types. On average, CD274 mutations are found in 0.96% of all cancers; the most common types are breast cancer, gastric cancer, lung cancer, colon cancer, bladder cancer, and prostate cancer [33].
- All cancers have genome instability as a hallmark. RMI2 is an important element of the BLM-TopoIIIa-RMI1-RMI2 complex that supports genome stability. Several studies have shown the upregulated expression of RMI2, which is caused tumor progression in cervical cancer, lung cancer, and prostate cancer [34].
- The MDS2 is a gene that encodes a protein that functions in the onset of myelodysplastic syndrome (MDS). Multiple mutations in MDS2 have been shown in breast and ovarian cancer. On average, MDS2 mutations are found in 0.09% of all cancers; the most common types are breast cancer, appendix cancer, lung cancer, and colon cancer [35].
- Tropomyosin-receptor kinase fused (TFG) encodes a protein which is a maintained regulator of protein secretion that controls the export of materials from the endoplasmic reticulum. TFG belongs to the systems that control cell size and is implicated in apoptosis and cell proliferation regulatory mechanisms. The TFG fusion proteins play a role in oncogenesis, with the activity of TFG fusion proteins promoting tumor development. Multiple mutations in TFG have been shown in intestinal cancer, lung cancer, and stomach cancer. On average, TFG mutations are found in 0.19% of all cancers; the most common types are breast cancer, colon cancer, and lung cancer [36].
- The U2AF1 encodes for a member of the spliceosome. This protein plays a vital role in RNA splicing. Multiple mutations in U2AF1 can cause irregular expression patterns of some genes affected in cancer pathogenesis. On average, U2AF1 mutations are found in 1.5% of all cancers; the most common types are acute myeloid leukemia, colon cancer, and lung cancer [30].
- The SRSF3 is a member Ser/Arg-rich (SR) proteins family. As a potential diagnostic and prognostic biomarker, SRSF3 is overexpressed in various types of cancer, including cancer of the breast, retinoblastoma, ovarian cancer, gastric cancer, head and neck cell squamous, colorectal cancer, cervical cancer and hepatocellular carcinoma (HCC). Recent studies also show SRSF3 upregulation in mesenchymal tumors [37].
- Previous studies showed that ATF1 plays a crucial role in carcinogenesis and participates in multiple cellular processes, including cell transformation, cell cycle, DNA damage, and apoptosis. ATF1 is overexpressed in various types of cancer, including lymphomas, nasopharyngeal carcinoma, and melanoma. However, other studies have shown that ATF1 acts as a tumor suppressor in breast and colorectal cancer [38].
- SDHC is a gene that encodes a protein as a part of succinate dehydrogenase. Multiple types of mutations in SDHC have been observed in ovarian cancer and pancreatic cancer. On average, SDHC mutations are found in 1.5% of all cancers; the most common types are lung cancer, breast cancer, pancreatic cancer, colon cancer, and bladder cancer [30].
- HOXD11 is a member of HOX family, which encodes transcription factors that control different physiological processes. Recent studies have shown that HOXD11 is involved in tumor development and helps control gene expression. Multiple types of mutations and changes in expression in HOXD11 have been observed in lung cancer, Oral Squamous Cell Carcinoma, prostate cancer, ovarian cancer, and Head and Neck Squamous Cell Carcinoma. HOXD11 may also change cell growth, clonality, and metastatic potential in Ewing sarcoma [39].
- The protein coded by the ZRSR2 gene plays a vital role in RNA splicing. Multiple types of mutations in ZRSR2 have been observed in chronic myelomonocytic leukemia and chronic lymphocytic leukemia. These mutations can drive abnormal expression patterns of some genes involved in cancer pathogenesis. On average, ZRSR2 mutations are found in 1.2% of all cancers; the most common types are lung cancer, breast cancer, colon cancer, and ovarian cancer [30].

#### 3.1.1 Signaling pathways associated with high score genes

One of the effective strategies for finding appropriate therapeutic approaches for cancer is identifying molecular pathways and specifying important genes in these pathways. Finding a new set of infrequently mutated genes with important properties can identify new pathways in different cancers and introduce them for further study. Therefore, we looked into the signaling pathways related to these genes and presented more information about them in Table1. We also studied the significant signaling pathways associated with our 200 top-selected mutated cancer-related genes. Table 2 shows some of the significant signaling pathways for these 200 top selected genes and the average Laplacian Scores of the genes for each of these pathways.

1. hsa04068: FoxO signaling pathway The first pathway with the highest score in Table 1 is the FoxO pathway. FoxO, as a family of transcription factors (FoxOs), has a direct role in cellular proliferation, oxidative stress response, and tumorigenesis. FoxOs are commonly inactivated by phosphorylation by several protein kinases such as AKT and PKB. The PI3K-Akt-FoxO signaling pathway has a significant role in various physiological processes such as cellular energy storage, growth, and survival [44]. One of the critical genes in this pathway that our algorithm has identified as a top gene is the transcriptional repressor factor CTCF. Recent studies show the effect of the CTCF factor on some cancers like prostate cancer by regulating the FoxO pathway [45]. The CTCF downregulates, or inhibition also governs the FoxO signal pathway and delays tumor growth. Therefore, the overexpression or genetic modification of CTCF affects the regulation of the FoxO pathway.

**Table 1.** Signaling pathways related to infrequently mutated genes.
2. hsa04015: MAPK signaling pathway The second pathway is Mitogen-Activated Protein Kinase (MAPK). The cascade of this pathway is a highly protected module that plays an essential role in different processes such as cell proliferation, differentiation, and migration, and any deviation from the precise control of this signaling pathway initiates many diseases [46], including various types of cancer. This signaling pathway has different signaling paths to the cell nucleus that the protein members of the MAPK /ERK chain (or Ras-Raf-MEK-ERK) are recognized by our algorithm. Studies show that the ERK signaling pathway plays a crucial role in tumorigenesis, migration, and invasion [47].
3. hsa04151: PI3K-Akt signaling pathway The phosphatidylinositol 3-kinase-Protein Kinase-B (PI3K-AKT) plays an important role in intracellular physiological regulation. Various oncogenes and growth factor receptors stimulate this signaling pathway, such as MET, KIT, EGFR, and ERBB3, which our algorithm recognizes. This signaling pathway also contains important genes such as PI3K, PTEN, mTOR, and JAK, which our algorithm recognizes. These gens induce cell proliferation, stem cell differentiation, and tumor suppressors in metabolic regulation. Disruption of this pathway and mutations in any of these genes can exhaust the cell of the natural process. This pathway is involved in cancer progression, and dysregulation of the PI3K pathway can be crucial in the cancer process [48].
4. hsa04014: Ras signaling pathway One of the critical signaling pathways in cellular activity is the Ras signaling pathway. Abnormal activation of Ras proteins (including RRAS2, MRas, HRas, KRas, and NRas) is the primary stimulus of oncogenes that has an essential role in the main signaling pathway in cancer. Mutations of Ras proteins such as KRas, which our method recognizes, cause cancer development. Meantime, the mutation in the regulatory ligands like EGFR and EGR, as other top mutated genes identified by our algorithm, cause the activation of their downstream signaling cascade [49].
5. hsa04012: ERBB signaling pathway The ERBB tyrosine kinase family members demonstrate some of the most generally changed proteins in cancer. Anomalous tyrosine kinase activation via gene alterations can cause tumorigenesis, tumor growth, and progression. This signaling pathway also contains important genes such as PI3K, CBLB, mTOR, and KRAS, which our algorithm recognizes. Oncogenic alterations of genes encoding members of the ERBB family, leading to unusual ERBB signaling and driving tumor growth, have been reported in different types of cancer, such as breast, lung, and gastrointestinal cancers. Recent studies show that the ERBB family’s signaling abnormalities and mutations are essential in escaping antitumor immunity in the cell process [50].
6. hsa04072: mTOR signaling pathway Mammalian target of rapamycin (mTOR) participates in multiple signaling pathways and controls cell proliferation, autophagy, and apoptosis. Studies show that the mTOR signaling pathway is related to different diseases, such as various types of cancer. This signaling pathway is often activated in tumors and plays an essential role in tumor metabolism. Therefore, the mTOR signaling pathway could effectively target through anti-tumor therapy studies [51].

| Gene name | Signaling pathway |
| --- | --- |
| FSTL3 | Ample FSTL3 expression promotes epithelial-mesenchymal transition (EMT) and improves aerobic glycolysis to positively affect cancer cells' invasive and metastatic capacity by activating the β-Catenin pathway. Results of [24] show that FSTL3 could be a bridging molecule in the crosstalk between HIPPO/YAP1 and Wnt/β-Catenin pathways and that FSTL3 is an essential regulatory factor of the β-Catenin molecular mechanisms in cancer. [24]. |
| COX6C | The expression level of COX6C was remarkably up-regulated in different cancers such as gastric and lung. It has been reported that overexpression of COX6C could promote the proliferation and decrease the apoptosis of cancer cells through activation of the oxidative phosphorylation pathway [25]. |
| SSX2 | It has been shown that the SSX proteins are activated in several critical mitogenic pathways, such as MAPK and Wnt [26]. |
| LMO1 | Studies show that LMO1 promoted the proliferation, aggression and migration of cancer cells by activation of NF-κB pathway [40]. |
| TNFRSF17 | Recent studies show that overexpression of TNFRSF17 in cells activates the MAPKs pathway, specifically JNK and p38 kinase, NF-κB, and Elk-1 [28]. |
| PD-L2 | Studies show the potential role of PD-L2 in regulating some pathways involved in cancer cell aggressiveness. They showed the modulation of ERK and Akt/PKB pathways are considered through PD-L2 [29]. |
| POU5F1 | Results of [31] demonstrate that IGF-IR/IRS-1/PI3K/AKT/GSK3β cascade-mediated regulation of POU5F1 and construction of β-catenin/POU5F1/SOX2 complex is essential for the retention of the self-renewal and tumorigenicity in cancer [31]. |
| HMGA1 | Recent studies show that HMGA1 is a critical transcription factor involved in multiple biological pathways, such as the TNF-α/NF-κB, EGFR, Hippo, Ras/ERK, Akt, Wnt/β-catenin and PI3-K/Akt pathways. In all of these pathways, HMGA1 targets various downstream genes [32]. |
| PD-L1 | PD-1/PD-L1 pathway regulates the induction and maintenance of immune tolerance within the tumor microenvironment. Recent studies show that PD-L1 is involved in multiple essential pathways, such as PI3K/AKT, MAPK, JAK/STAT, WNT, and NF-κB pathways [33]. |
| RMI2 | Results of KEGG enrichment analysis indicated that RMI2 was significantly associated with the p53 signaling pathway [34]. |
| MDS2 | Recent studies show that knockdown of SPAG6 significantly increased the apoptosis of MDS cells by inducing the activation of tumor suppressor genes, such as p53 and PTEN. SPAG6 knockdown induced autophagy via the AMPK/mTOR/ULK1 signaling pathway in MDS2 cells, and inhibiting autophagy decreased SPAG6 knockdown-mediated apoptosis [35]. |
| TFG | Recent studies show that TFG is involved in the NF-κB and MAPK pathways, and activation of MAPK pathway occurs in various cancers, and the NF-κB pathway is essential in inhibition of apoptosis and treatment resistance in cancers [36]. |
| U2AF1 | The KEGG pathway enrichment analysis results showed that U2AF1 was involved in several biological pathways, such as FoxO and PI3K/Akt signaling pathways [41]. |
| SRSF3 | Recent studies show that SRSF3 as an oncogene manipulates various cell functions by regulating many pathways, such as p53, JNK, Ras, Wnt, and HER2 signaling pathways [37]. |
| ATF1 | Recent studies showed that ATF1 activates a subset of genes related to apoptosis, Wnt, TGF-β, and MAPK pathways, and these consequences could increase the risk of various cancers [?]. |
| SDHC | Recent studies showed that SDH activity has a significant role in regulating oncogenic signaling pathways, such as those associated with NF-κB [42]. |
| HOXD11 | Recent studies showed that HOXD11 is involved in various cancer-related signaling pathways such as cell cycle, DNA replication, ECM receptor interaction, and focal adhesion [39]. |
| ZRSR2 | Recent studies showed that ZRSR2 is involved in various cancer-related signaling pathways, such as TLR signaling pathway [43]. |

Figure 3 shows the cross-talk between all of these significant signaling pathways in different types of cancer.

**Fig 3.**
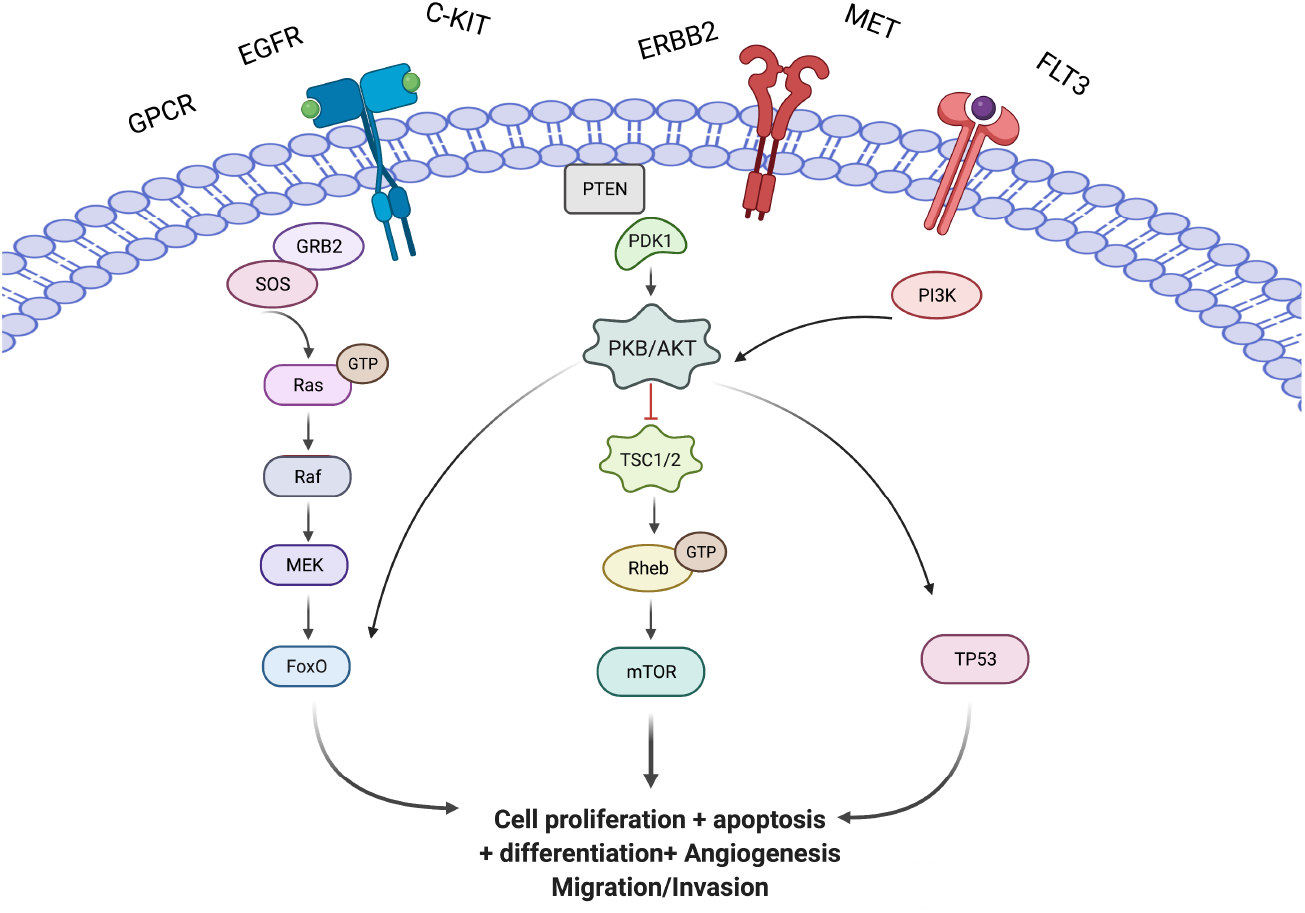
Cross-talk between mentioned signaling pathways in different types of cancer

### 3.2 Evaluation of the proposed clusters based on Gene Ontology

To gain a better understanding of the biological function and physical interaction of the genes in each cluster, we have performed an analysis of the GO term annotations of the obtained clusters from our method with the help of the Database for Annotation, Visualization, and Integrated Discovery (DAVID) [52]. Figure 4 shows significant GO terms for each cluster resulting from DAVID.

**Fig 4.**
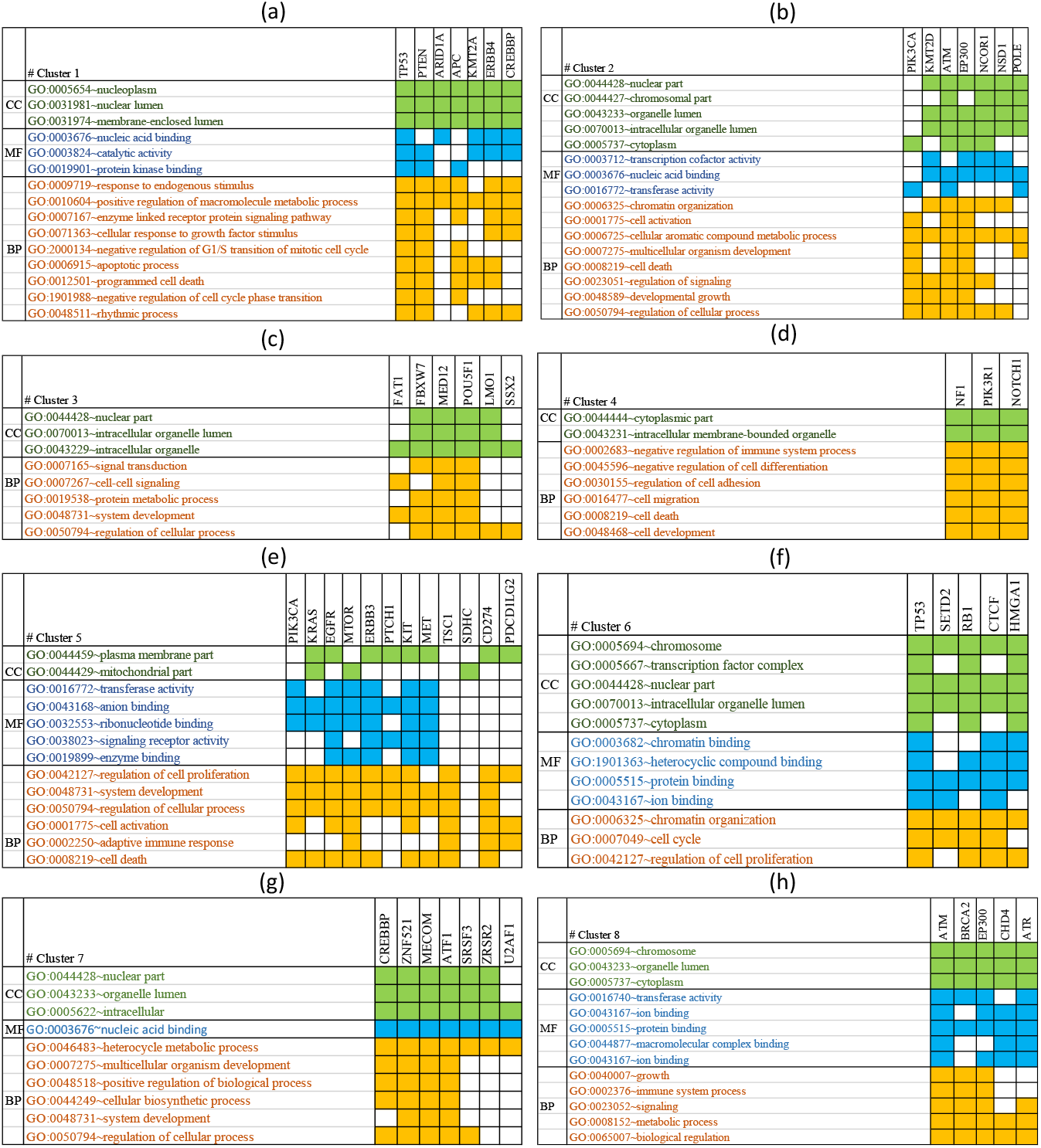
Significant GO terms for each cluster.

The first cluster, the most important cluster, is obtained by expanding the TP53 gene with an average Laplacian Score of 360.58 for its genes, which contains TP53, PTEN, ARID1A, APC, KMT2A, ERBB4, and CREBBP genes. The accumulation of these genes is higher in the nucleoplasm and nuclear lumen region. These genes also participate in various functions, including binding to nucleotide acid and catalytic activity. From the above genes, genes such as ATP, PTEN, and TP53 have the function of binding to protein kinases. These genes are active in many biological processes, some mentioned in Figure 1 (a), including GO:0006915 ∼ apoptotic process and GO:0012501 ∼ programmed cell death.

The second cluster is obtained by expanding the PIK3CA gene with an average Laplacian Score of 320.75 for its genes, which contain PK3CA, KMT2D, ATM, EP300, NCOR1, NSD1, and POLE genes. Most of these genes are concentrated in the intracellular organelle lumen or cytoplasm, and all of them have an activity of GO:0003676 ∼ nucleic acid-binding. These genes are involved in critical biological processes such as GO:0001775 ∼ cell activation, GO:0048589 ∼ developmental growth, and GO:0008219 ~ cell death. Figure 1 (b) shows significant GO terms associated with the genes of this cluster.

The third cluster is obtained by expanding the FAT1 gene with an average Laplacian Score of 315.48 for its genes, which contains FAT1, FBXW7, MED12, POUF1, LMO1, and SSX2 genes. All genes in this cluster are known as regulators of biological processes and play a role in signal transmission. Figure 1 (c) shows significant GO terms associated with the genes of this cluster.

The fourth cluster, the smallest cluster, is obtained by expanding the NF1 gene with an average Laplacian Score of 301.88 for its genes, which contains NF1, PIK3R1, and NOTCH1 genes. The accumulation of these genes is cytoplasm and an intracellular membrane-bounded organelle. These genes are also involved in critical biological processes such as GO:0016477∼ cell migration, GO:0048468 ~ cell development, and GO:0008219 ∼ cell death. Figure 1 (d) shows significant GO terms associated with the genes of this cluster.

The fifth cluster, the largest cluster is obtained by expanding the KRAS gene with an average Laplacian Score of 286.72 for its genes, which contains PIK3CA, KRAS, EGFR, mTOR, ERBB3, PTCH1, KIT, MET, TSC1, CD274, PDCD1LG2 genes. These genes are involved in critical biological processes such as GO:0002250 ∼ adaptive immune response, GO:0008219 ∼ cell death, and GO:0042127 ∼ regulation of cell proliferation. Figure 1 (e) shows significant GO terms associated with the genes of this cluster.

The sixth cluster is obtained by expanding the SETD2 gene with an average Laplacian Score of 292.53 for its genes, which contains TP53, SETD2, RB1, CTCF, and HMGA1 genes. These genes are involved in critical biological processes such as GO:0042127 ∼ regulation of cell proliferation and GO:0007049 ∼ cell cycle. Figure 1 (f) shows significant GO terms associated with the genes of this cluster.

The seventh cluster is obtained by expanding the CREBBP gene with an average Laplacian Score of 284.07 for its genes, which contain CREBBP, ZNF521, MECOM, ATF1, SRSF3, ZRSR2, and U2AF1 genes. In this cluster, some genes such as ATF1, SRSF3, and ZRSR2, have fewer mutations than other genes. These genes are involved in critical biological processes such as GO:0048518 ∼ positive regulation of biological process and GO:0048731 ∼ system development. Figure 1 (g) shows significant GO terms associated with the genes of this cluster.

The last cluster is obtained by expanding the BRCA2 gene with an average Laplacian Score of 300.52 for its genes, which contain ATM, BRCA2, EP300, CHD4, and ATR genes. The accumulation of these genes is more in the chromosome and cytoplasm regions and they are involved in critical biological processes such as GO:0040007 ∼ growth and GO:0002376 ∼ immune system process. Figure 1 (h) shows significant GO terms associated with the genes of this cluster.

### 3.3 Evaluation of the proposed mutated modules

In the previous subsection, we evaluated each of the proposed clusters. We showed that each cluster participates in important biological processes such as cell proliferation, migration, and cell growth. In this subsection, we evaluated the result of our method to find the important modules for each cancer. We studied the genes in each cluster that simultaneously have several mutations in multiple cases to find driver modules. Table 3 shows the set of modulated genes for each cancer. The first column of Table 3 shows the module number corresponding to each cluster. The second column shows the corresponding cluster number, and the third and fourth columns show the genes found in more than 10% and less than 10% of the cases simultaneously. For example, among 1379 patients with breast cancer, 65 patients in cluster 2 had mutations in the NCOR1 gene. Of these 65 patients, 38 patients had mutations in NCOR1 and PIK3CA genes simultaneously. Therefore, we have reported these two genes as a driver module of breast cancer. The important point to finding the module in this section is that, like previous studies, we have examined the number of simultaneous mutations in the most significant number of cases. Genes with fewer mutations are expected to participate in fewer modules. Therefore, if we want to see genes with fewer mutations in our modules, we should define other criteria than the number of mutations. Figure 5 shows the number of cases in the modules introduced by our algorithm in each cancer separately. For example, corpus uteri cancer has modules in all clusters, and modules 3, 4, and 8 are found in more cases than the other five modules.

**Table 2.** Significant signaling pathways with the high average LS.

| Signaling pathway | Ave. LS | No. gene |
| --- | --- | --- |
| hsa04068: FoxO signaling pathway | 314.4 | 10 |
| hsa04010: MAPK signaling pathway | 308.8 | 15 |
| hsa04151: PI3K-Akt signaling pathway | 302.3 | 20 |
| hsa04014: Ras signaling pathway | 299.7 | 16 |
| hsa04012: ERBB signaling pathway | 295 | 10 |
| hsa04072: mTOR signaling pathway | 290.8 | 9 |

**Table 3.** A set of obtained important modules for each cancer separately.

| Cancer type | No. Modules | No. Clusters | Genes Mutated in > 10% cases | Genes Mutated in < 10% cases |
| --- | --- | --- | --- | --- |
| Bladder | Module 1 | Cluster 1 | TP53, PTEN | ARID1A |
|  | Module 2 | Cluster 2 | KMT2D, EP300 |  |
|  | Module 5 | Cluster 5 |  | PIK3CA, ERBB3 |
|  | Module 6 | Cluster 6 | TP53, RB1 |  |
|  | Module 8 | Cluster 8 | EP300 | ATM |
| Brain | Module 1 | Cluster 1 | TP53 | PTEN |
|  | Module 2 | Cluster 2 | KMT2D | POLE |
|  | Module 3 | Cluster 3 |  | FAT1, FBXW7 |
|  | Module 4 | Cluster 4 | NF1, PIK3R1 |  |
|  | Module 5 | Cluster 5 | EGFR | PIK3CA |
|  | Module 6 | Cluster 6 | SETD2, RB1 | HMGA1 |
| Breast | Module 1 | Cluster 1 | TP53, PTEN | ARID1A |
|  | Module 2 | Cluster 2 | PIK3CA | NCOR1 |
|  | Module 4 | Cluster 4 |  | NF1, PIK3R1 |
|  | Module 5 | Cluster 5 | PIK3CA | KRAS, |
|  | Module 6 | Cluster 6 | TP53 | SETD2 |
| Lung | Module 1 | Cluster 1 | TP53, PTEN, ARID1A |  |
|  | Module 2 | Cluster 2 | KMT2D, ATM, |  |
|  | Module 3 | Cluster 3 | FAT1, FBXW7 |  |
|  | Module 4 | Cluster 4 | NF1, NOTCH1 |  |
|  | Module 5 | Cluster 5 | KRAS, EGFR | SDHC |
|  | Module 6 | Cluster 6 | TP53, RB1 |  |
|  | Module 8 | Cluster 8 | ATM | EP300 |
| Cervix uteri | Module 1 | Cluster 1 |  | PTEN, ARID1A |
|  | Module 2 | Cluster 2 |  | PIK3CA, EP300 |
|  | Module 3 | Cluster 3 |  | FAT1, FBXW7 |
| Colon | Module 1 | Cluster 1 | TP53, PTEN |  |
|  | Module 2 | Cluster 2 | POLE | KMT2D |
|  | Module 5 | Cluster 5 | KRAS | PIK3CA, |
|  | Module 6 | Cluster 6 | SETD2 | TP53 |
|  | Module 7 | Cluster 7 | CREBBP | ATF1 |
|  | Module 8 | Cluster 8 | ATM, EP300, CHD4 |  |
| Corpus uteri | Module 1 | Cluster 1 | PTEN, ARID1A, KMT2A | TP53, ERBB4 |
|  | Module 2 | Cluster 2 | PIK3CA, KMT2D, NSD1, POLE | ATM |
|  | Module 3 | Cluster 3 | FAT1, FBXW7 |  |
|  | Module 4 | Cluster 4 | NF1, PIK3CA |  |
|  | Module 5 | Cluster 5 | PIK3CA, mTOR, ERBB3, PTCH1 | EGFR, KIT, MET, TSC1 |
|  | Module 6 | Cluster 6 | SETD2, RB1, CTCF | TP53 |
|  | Module 7 | Cluster 7 | CREBBP | ATF1 |
|  | Module 8 | Cluster 8 | CHD4 | ATM |
| Esophagus | Module 1 | Cluster 1 |  | TP53, ARID1A |
|  | Module 2 | Cluster 2 | KMT2D, ATM | PIK3CA |
|  | Module 5 | Cluster 5 | KRAS | PIK3CA |
| Kidney | Module 1 | Cluster 1 |  | TP53, PTEN |
| Liver | Module 1 | Cluster 1 |  | TP53, ARID1A |
|  | Module 6 | Cluster 6 |  | TP53, SETD2 |
| Ovary | Module 1 | Cluster 1 | TP53, KMT2A |  |
|  | Module 6 | Cluster 6 | TP53 | SETD2 |
| Pancreas | Module 1 | Cluster 1 | TP53 | ARID1A |
|  | Module 5 | Cluster 5 | KRAS | PIK3CA |
| Skin | Module 1 | Cluster 1 |  | TP53, PTEN |
|  | Module 2 | Cluster 2 | KMT2D, ATM |  |
|  | Module 3 | Cluster 3 | POUSF1, SSX2 |  |
|  | Module 6 | Cluster 6 |  | TP53, SETD2 |
| Stomach | Module 1 | Cluster 1 | TP53, PTEN | ARID1A, ERBB4 |
|  | Module 2 | Cluster 2 |  | PIK3CA, KMT2D, ATM |
|  | Module 4 | Cluster 4 |  | NF1, PIK3R1 |
|  | Module 5 | Cluster 5 |  | PIK3CA, KRAS, TSC1 |
|  | Module 7 | Cluster 7 |  | CREBBP, ZNF512 |
|  | Module 8 | Cluster 8 | ATM, CHD4, ART |  |
| Uterus | Module 1 | Cluster 1 |  | TP53, PTEN |
|  | Module 2 | Cluster 2 |  | PIK3CA, KMT2D |
|  | Module 7 | Cluster 7 | ZNF512 | ZRSR2 |

**Fig 5.**
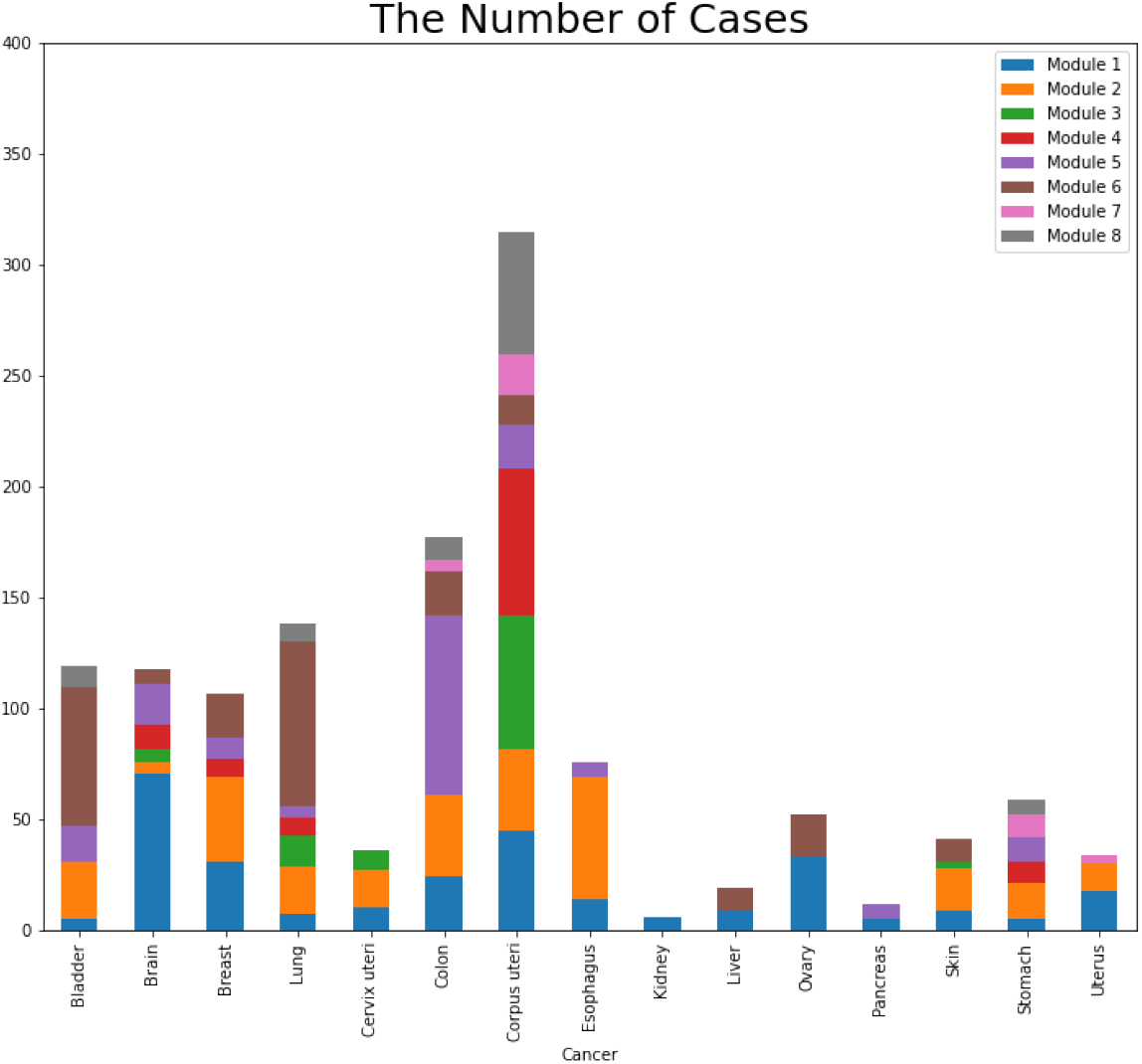
The number of cases in each module in each cancer separately.

## Conclusion and Discussion

New sequencing technologies and improving genomics data help us identify cancer-related genes and modules in various cancers. Most previous studies focus on using statistical methods to identify high-frequency mutation genes. Finding these mutation genes is important in determining the cancer progression mechanism. The critical point is that some critical genes do not have high mutation frequencies and can not be identified depending on the number of mutations and statistical techniques. In this study, we used a machine learning method to find important cancer genes with low-frequency mutations along with the driver genes with high-frequency mutations. For this purpose, we presented a novel three-step method to identify driver gene sets with mutually exclusive mutations.

In the first step, we extracted 576 cancer-related genes for 15 common cancers reported on TCGA and constructed a weighted graph for the corresponding mutations of these genes. The weight of the associated edge between two genes in this network is based on the number of common cases that contain these mutated genes simultaneously. Since the problem of finding candidate driver genes is still an open question, it can be studied as a problem without an exact answer. We used an unsupervised learning method to determine an efficient set of mutated cancer genes to find an appropriate response to this question. We defined six informative features for each gene and calculated the score for these features with the help of the mentioned unsupervised machine learning method for each gene. Afterward, we introduced 200 high-score genes with meaningful relationships to cancer as candidate genes for more investigation (Figure 2). Our method proposed some genes, such as TP53, FAT4, and KMT2C, with high-frequency mutations as high-score genes that are presented through other statistical methods. In addition to these genes, our method also identified some genes, such as FSTL3, SSX2, and MDS2, with low-frequency mutations. We briefly studied these genes with low-frequency mutations and examined the association of each of these genes with different types of cancer. In addition, we studied the KEGG signaling pathways of the set of high-score genes. We also examined the roles of these high-score genes and the effects of mutations and abnormalities of these genes in the proposed set of signaling pathways in the different cellular processes such as proliferation and migration.

Genomic analysis of different types of mutation in genes indicates the mutation heterogeneity problem. Genes should be accepted as a module rather than as individuals in order to solve this heterogeneity issue. We used the knowledge of gene binding in protein-protein interaction networks and the information on the biological processes of each of these genes to detect the high-score genes and identify cancer-stimulating modules with high accuracy. For this purpose, we created a network based on information about the physical interactions of genes and the biological processes of these genes. We added weight to each node of this network with the help of the Laplacian Score. Then, we proposed a heuristic algorithm and clustered the network. We introduced 8 top clusters with the highest Laplacian Score as cancer clusters from these clusters. To better understand the biological function of the genes in each cluster, we analyzed the GO term annotations of the genes for each cluster with the help of the DAVID tool. Finally, we studied the genes in each cluster that simultaneously have several mutations in multiple cases to find driver modules for each cancer separately.

It can be concluded that the methods that filtrate mutation data based on the most mutated genes and pre-defined network modules may lose important information about genes with a lower frequency of mutations. The mutual exclusivity signal may be biased toward recognizing gene sets that have a large proportion of their coverage in highly mutated genes. Although cancer-related genes have been shown to be involved in numerous pathways, few methods have been developed to identify the candidate driver gene sets with different mutation frequencies. We proposed a method to detect candidate driver genes with varying mutation frequencies and showed their critical role in cancer progression.

